# Co-Component Signal Transduction Systems: fast-evolving virulence regulation cassettes discovered in enteric bacteria

**DOI:** 10.1101/2022.04.08.487670

**Authors:** Lisa N. Kinch, Qian Cong, Jananee Jaishankar, Kim Orth

**Affiliations:** Department of Molecular Biology, University of Texas Southwestern Medical Center, Dallas, TX 75390, USA; Howard Hughes Medical Institute, University of Texas Southwestern Medical Center, Dallas, TX 75390, USA; Department of Eugene McDermott Center for Human Growth & Development, University of Texas Southwestern Medical Center, Dallas, TX 75390, USA; Department of Biophysics, University of Texas Southwestern Medical Center, Dallas, TX 75390, USA; Department of Biochemistry, University of Texas Southwestern Medical Center, Dallas, TX 75390, USA

**Keywords:** co-component signal transduction system, virulence transcription regulation, enteric bacteria, protein structure prediction, protein sequence evolution

## Abstract

Bacterial signal transduction systems sense changes in the environment and transmit these signals to control cellular responses. The simplest one-component signal transduction systems include an input sensor domain and an output response domain encoded in a single protein chain. Alternately, two-component signal transduction systems transmit signals by phosphorelay between input and output domains from separate proteins. The membrane tethered periplasmic bile acid sensor that activates the *Vibrio parahaemolyticus* type III secretion system adopts an obligate heterodimer of two proteins encoded by partially overlapping *VtrA* and *VtrC* genes. This co-component signal transduction system binds bile acid using a lipocalin-like domain in VtrC and transmits the signal through the membrane to a cytoplasmic DNA-binding transcription factor in VtrA. Using the domain and operon organization of VtrA/VtrC, we identify a fast-evolving superfamily of co-component systems in enteric bacteria. Accurate machine learning-based fold predictions for the candidate co-components support their homology in the twilight zone of rapidly evolving sequence and provide mechanistic hypotheses about previously unrecognized lipid-sensing functions.

**Significance statement:** Using the domain and operon organization of VtrA/VtrC, combined with fold predictions, we identify new co-component signal transduction systems in enteric bacteria that likely regulate virulence. We observe that the heterodimeric VtrA/VtrC periplasmic bile acid receptor controlling *Vibrio parahaemolyticus* T3SS2 is a distant homolog of the ToxR/ToxS master regulator of virulence and has evolved beyond confident sequence recognition. Exploiting the newly developed machine learning methods for structure prediction, we observe a VtrC-like lipocalin fold for both the ToxS periplasmic domain and for other detected periplasmic sensor components. This structure prediction supports the divergent evolution of VtrA/VtrC-like co-component signal transduction systems and suggests a role for lipid sensing in regulating virulence in enteric bacteria.

## Introduction

Bacterial transmembrane signal transduction systems use periplasmic proteins as inputs for sensing changes in extracellular cues. The input sensors control a variety of adaptive cellular responses by stimulating different intracellular signaling proteins, including chemoreceptors, sensor kinases, and diguanylate cyclases/phosphodiesterases, among others(1). For example, the well-studied Tar and Trg chemoreceptors from *E. coli* transmit environmental signals through a cytoplasmic CheA sensor histidine kinase and CheY receiver as part of a two-component system. These chemoreceptors use a periplasmic ligand binding domain to interact with maltose-, ribose- and galactose-binding proteins, which mediates chemotaxis towards these sugars(2). Often, genes that encode periplasmic solute binding proteins that stimulate sensor histidine kinases are found near their signaling genes (1). Some two-component systems have periplasmic solute binding domains fused directly to their intracellular signaling domains. The virulence regulatory PhoP/PhoQ two-component system includes a PhoQ periplasmic sensor domain connected by a transmembrane helix (TMH) to a cytoplasmic histidine kinase (Fig. 1*A*). Under low periplasmic Mg^2+^ (or Ca^2+^) conditions, PhoQ signals to the receiver domain of its DNA-binding transcriptional regulator PhoP to activate transcription. The PhoP and PhoQ encoding genes are also neighboring(3, 4).

**Figure 1.**
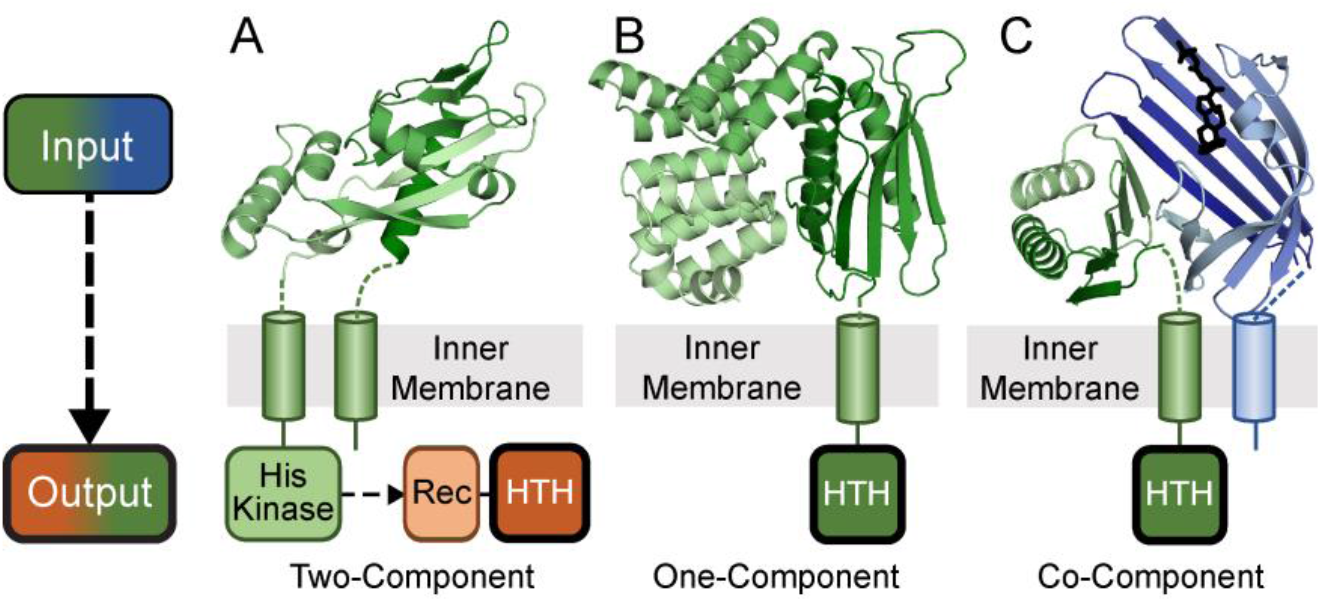
Three transmembrane signaling systems found in enteric bacteria. Extracellular inputs (above, cartoon color scale from dark N-terminus to light C-terminus) and cytoplasmic outputs (below) are depicted for three representative types of bacterial signaling systems: **A)** Two-component PhoQ (PDB: 1yax) periplasmic input sensor domain (green) is attached through a TMH to an intracellular histidine kinase that signals to a second component: the PhoP output response regulator. **B)** One-component CadC (PDB:3ly7) periplasmic domain (green) is attached through a TMH to the output HTH domain. **C)** Co-component VtrA/VtrC (PDB: 5kew) periplasmic input domains (green/blue), with VtrC attached through a TMH to the output HTH domain.

One-component signal transduction systems typically include an input domain that senses environmental stimuli fused to an output domain that controls an adaptive response. One-component regulators include many of the same input and output domains found in their two-component system counterparts, yet they lack the histidine kinase and receiver(5-7). The diverse domain repertoire of one-component bacterial signal transduction systems most commonly includes various intracellular small molecule-binding input modules that stimulate DNA-binding helix turn helix (HTH) output modules to regulate gene expression(6). However, a small subset of one-component transcription factors possesses TMHs and can respond to extracellular stimuli(6, 8, 9). The integral membrane transcription factor CadC exemplifies this type of one-component system, with a periplasmic sensor fused to a cytoplasmic helix-turn-helix (HTH) DNA-binding domain (Fig. 1*B*). The periplasmic CadC input module responds directly to low pH to activate the expression of the *CadBA* operon and maintain a neutral cytoplasmic pH. The CadC sensor also binds the feedback inhibitor cadaverine to shut off the system(9-12). The input modules from many of these signaling systems are diverse, lack easily detected sequence similarity to known domains and can require co-regulators for activity(13, 14).

The membrane tethered ToxR transcription factor that activates the *Vibrio cholerae* virulence regulatory cascade has long been described as a one-component system(15-17). The domain architecture of ToxR includes an N-terminal DNA-binding HTH, a TMH and a periplasmic domain with a recently determined structure(18, 19). The ToxR HTH belongs to the CadC superfamily of transcription factor DNA-binding domains, while the periplasmic sensor combines an N-terminal αβ-sandwich with a C-terminal α-helical ARM repeat(9, 11). However, the ToxR periplasmic domain structure lacks the CadC sensor subdomain organization and instead requires a second unrelated periplasmic protein ToxS for activity(20). The ToxS co-regulator is encoded by a neighboring gene from the ToxR operon (20, 21), and the two gene products form a complex that stabilizes ToxR and renders it resistant to proteolytic cleavage(18, 22). Thus, instead of ToxR serving as a traditionally recognized one-component system, we suggest the periplasmic ToxR/ToxS heterodimer serves as a sensor in a “co-component” signal transduction system. The ToxR/ToxS sensor responds to various environmental cues, such as bile salts, pH and redox state(18, 19, 23-25), although the molecular mechanism of activation by these factors is not fully understood.

The ToxR CadC-like HTH family includes another membrane-tethered transcription factor (VtrA) from *Vibrio parahaemolyticus* (*V. parahaemolyticus*). VtrA possesses a similar domain composition as ToxR and functions together with a neighboring genomic co-regulator (VtrC)(8). VtrA/VtrC serves as a bile salt sensor that ultimately activates transcription of the Type III Secretion System 2 (T3SS2) virulence system during *V. parahaemolyticus* infection of the human gut(13, 26). The periplasmic VtrA/VtrC structure adopts an obligate heterodimer that binds bile acid using a lipid-binding lipocalin-like fold from VtrC (13). Despite a significant divergence between the VtrA and ToxR periplasmic sequence, the two adopt a similar fold that defines the VtrA/ToxR periplasmic input modulator superfamily(13, 19). Given the fast-evolving nature of this input modulator superfamily, we suspect the corresponding ToxS could also be homologous to VtrC and adopt a lipocalin-like fold.

Protein folds can often be inferred from their evolutionary relationships to experimentally determined structures. These relationships have traditionally been detected at the sequence level by homology-based protein structure prediction methods. However, for fast evolving domains such as in the periplasmic regions of VtrA/VtrC and ToxR/ToxS, identifying homologs using sequence has proven difficult, even for the most sensitive methods(27-29). Here, we take advantage of the VtrA/ToxR transmembrane-containing domain organization and its preserved neighboring gene arrangement to identify candidate co-component systems in enteric bacteria that have evolved beyond sequence recognition. The structures of similar fast evolving sequences were recently predicted with accuracy approaching that of experimental structures by machine learning (30-33). Using these structure prediction methods(31, 32, 34), we provide evidence the identified candidates adopt VtrA/VtrC-like folds. The fold predictions suggest the two-gene cassettes encoding co-component signaling systems evolved from a common ancestor and provide functional implications for the new superfamily members (Fig. 1*C*). This example highlights the role of AI-based structure prediction in shifting a structure biology paradigm, where *de novo* predictions can now inform homology beyond sequence detection limits.

## Results and Discussion

### The *ToxR/ToxS* gene cassette encodes fast-evolving homologs of VtrA/VtrC

The periplasmic VtrA/VtrC structure adopts an obligate heterodimer that senses the bile acid taurodeoxycholate (TDC) using a lipocalin-like fold adopted by the VtrC component (Fig. 2*A*). The bile acid input transduces a signal through the membrane to the VtrA HTH DNA-binding domain and initiates transcription of the *V. parahaemolyticus* T3SS2 virulence secretion machinery. The two genes encoding the VtrA/VtrC co-component signal transduction system are neighboring in the *V. parahaemolyticus* genome and form an operon (Fig. 2*B*). Functional similarities between the VtrA/VtrC and ToxR/ToxS systems, and their preserved operon arrangement (Fig. 2*B*) suggest their homologous DNA-binding output domain relationship could extend across the whole two gene cassette. Accordingly, the periplasmic folds of VtrA and ToxR are related, adopting a similar αβ-sandwich fold (Fig. 2*C*). Despite this structure similarity, sensitive sequence detection methods starting from either of the periplasmic domain sequences confidently identify sequences from the same family, but not across families (SI Appendix, Table S1). Thus, VtrA and ToxR exemplify homologous transcription factors whose periplasmic input domains have evolved beyond sequence recognition.

**Figure 2.**
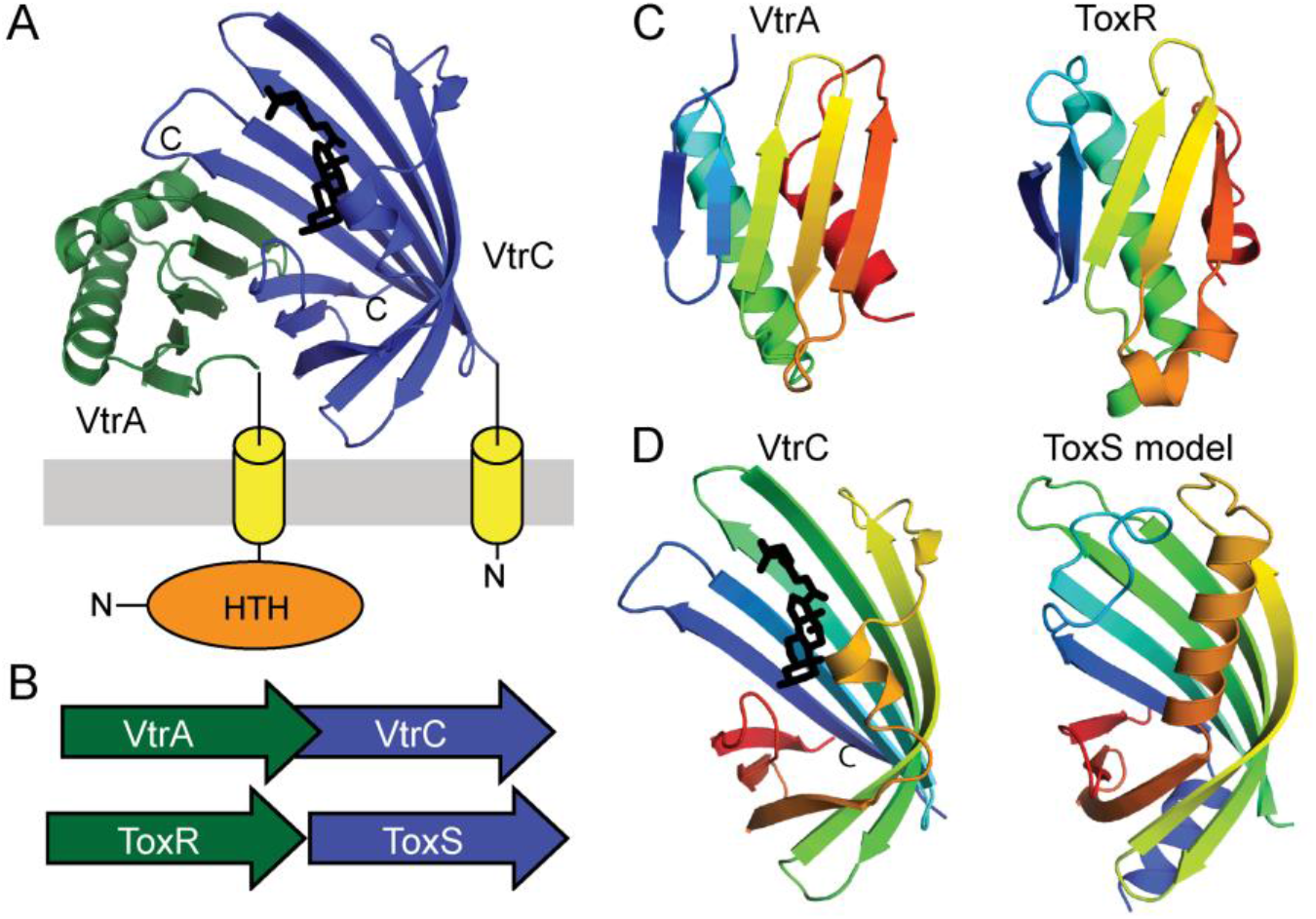
VtrA/VtrC T3SS2 virulence regulon is homologous to ToxR/ToxS. **A)** The VtrA/VtrC structure (PDB: 5kew) depicted in cartoon adopts an obligate heterodimer of the periplasmic VtrA (green) and VtrC (blue) that transmits an input signal from VtrC bound bile acid (black stick) through the membrane (gray) via the TMH (yellow cylinder) attached to the N-terminal VtrA DNA-binding HTH (orange sphere). **B)** The *VtrC* gene (green arrow) encoding the coregulator is downstream from the *VtrA* gene (blue arrow) encoding the transcription factor in an overlapping operon that is arranged similarly to the genes of the *ToxR* (green arrow) *ToxS* (blue arrow) operon. **C)** The labeled VtrA periplasmic domain colored in rainbow from the N-terminus (blue) to the C-terminus (red), adopts the same fold as the labeled ToxR periplasmic domain (PDB: 6utc). **D)** The labeled VtrC periplasmic domain that binds the bile acid TDC (black stick) adopts the same fold as the predicted ToxS structure model.

The rapidly diverging VtrA and ToxR periplasmic domains interact with their co-transcribed operon partners VtrC and ToxS, respectively. The VtrC structure adopts a lipocalin-like fold that typically binds lipid (Fig. 2*D*). However, the structure of the corresponding ToxR operon partner ToxS is unknown, and sensitive sequence detection methods fail to identify a relationship between ToxS and VtrC (SI Appendix, Table S1). Despite this lack of detectable sequence relationship, the ToxS sequence does identify an alternate lipocalin-like fold with borderline confidence and coverage (SI Appendix, Table S1). This hint at ToxS adopting a similar fold as its VtrC counterpart prompted us to build a structure model using newly developed structure prediction methods, including AlphaFold and ColabFold (31, 32, 34, 35). Indeed, the ToxS model adopts a lipocalin-like fold similar to VtrC (Fig 2*D*). High confidence is assigned to all residues comprising the core lipocalin-like domain. Together with the similar domain organization and periplasmic folds of ToxR and VtrA, this ToxS structure prediction supports the proposed homologous relationship between the *ToxR/ToxS* and *VtrA/VtrC* gene cassettes.

### Genome neighborhoods facilitate discovery of candidate VtrA/VtrC-like co-component systems

Given the divergent nature of the VtrA/VtrC periplasmic input components, we reasoned additional membrane tethered co-components might exist. Accordingly, sequence search using the VtrC periplasmic domain identified the PsaF superfamily with borderline confidence (SI Appendix, Table S1). PsaF works together with its neighboring membrane tethered transcription factor PsaE to control expression of the pH6 antigen adhesin virulence factor in *Yersinia* (36). The PsaE DNA-binding domain is related to both ToxR and another positive regulator of *Vibrio cholerae* virulence TcpP, which works together with its neighboring TcpH periplasmic input domain (37). Each of the transcription factors has the same domain organization, with an N-terminal helix-turn-helix (HTH) domain, followed by a single TMH, and a C-terminal periplasmic domain of unknown structure. Thus, the similar domain organization, transcription regulatory function, and neighboring gene organization suggest the VtrA/VtrC-like superfamily could exist as a mobile two-gene cassette that controls virulence in *Vibrio* as well as in other enteric pathogens.

While the periplasmic components of VtrA/VtrC, ToxR/ToxS, and PsaE/PsaF have diverged beyond confident sequence recognition, the CadC-like HTH retains significant sequence similarity and can be used to search for candidate co-component systems in other pathogenic bacteria. Starting from the VtrA and ToxR, we searched a database of well-studied bacterial pathogens with complete genomes of high sequence and annotation quality (SI Appendix, Table S2, 18 genomes). The VtrA/ToxR sequence search identified numerous CadC-like HTH-containing proteins from all represented genomes (252 sequences total). These identified sequences were filtered using TMH prediction, resulting in 28 transmembrane-tethered transcription factors from eight species of enterobacteria. To further filter these identified transcription factors for candidate VtrA/VtrC-like co-component signal transduction systems, their gene neighborhood was inspected for a similar potential co-translated operon organization. Adjacent downstream genes in the same orientation as the identified transcription factors were subjected to transmembrane and signal peptide prediction. One notable candidate co-component system from *Yersinia pseudotuberculosis* required an N-terminal extension of the predicted start site to include the TMH (BZ17_3565, for simplicity “BZ17_” is replaced with “YP” in the subsequent text). This alternate start site was likely miss-annotated due to its reading frame overlap with the upstream transcription factor (as was seen with the VtrA/VtrC genes) and is extended correctly in close orthologs from other *Yersinia pseudotuberculosis* strains.

Inspection of genomic neighborhoods excluded membrane tethered CadC one-component transcription factors, for which the VtrA HTH domain is named, as well as dismissed other pairs that lack input domains or neighboring proteins predicted to be in the periplasm. In addition to the known *Vibrio ToxR/ToxS, VtrA/VtrC*, and *TcpP/TcpH* gene cassettes, several gene pairs from other enteric bacteria passed the search criteria (Table 1). *Yersinia pseudotuberculosis* includes two novel gene cassette pairs of unknown function in addition to *TcpP/TcpH. Shigella flexneri* encodes a single candidate co-component pair of unknown function, while *Salmonella typhimurium* encodes three candidate pairs. Two are neighboring in the genome (STM0342 and STM0345), while the last represents the MarT (for membrane-associated regulator(38))/FidL cassette from pathogenicity island 3 (Spi-3). The *MarT/FidL* gene products are co-transcribed and activate expression of the MisL autotransporter, which functions as a host specific intestinal colonization factor(38-40). The noninvasive *E. coli* K12 strain includes a single candidate cassette *YqeI/YqeJ*. The *YqeI* and *YqeJ* genes are described as a remnant of the ETT2 (type III secretion system) pathogenicity island in K12 that is fully present in pathogenic strains like *E. coli* O157:H7 saki, which also encodes YqeI/YqeJ. In addition to this virulence regulator, the pathogenic *E. coli* O157:H7 saki strain encodes an additional cassette: *GrvA* (for Global Regulator of Virulence A(41))/*FidL*. The GrvA transcription factor activates the locus of enterocyte effacement (LEE)-dependent adherence.

**Table 1.** Candidate Co-Component Systems.

| Bacteria | ToxR ORF;Gene | Neighbor | Comment |
| --- | --- | --- | --- |
| *Vibrio parahaemolyticus* | VP0820;ToxR | ToxS | ToxS Lipocalin |
| *Vibrio parahaemolyticus* | VPA1332;VtrA | VtrC* | known |
| *Vibrio cholerae* | MS6_0632;TcpP | MS6_0633* | TcpH Lipocalin |
| *Vibrio cholerae* | MS6_0778;ToxR | MS6_0777 | ToxS Lipocalin |
| *Yersinia pseudotuberculosis* | BZ17_3564 | BZ17_3565* | Lipocalin |
| *Yersinia pseudotuberculosis* | BZ17_1189;PsaE | PsaF* | PsaF lipocalin |
| *Yersinia pseudotuberculosis* | BZ17_3283 | BZ17_3282* | Lipocalin |
| *Escherichia coli* K-12 | ECK2845;Yqel | ECK2846 | YqeJ Lipocalin |
| *Escherichia coli* Sakai | ECs_3704;Yqel | ECs_3705 | YqeJ Lipocalin |
| *Escherichia coli* Sakai | ECs_1274;GrvA | ECs_1273 | FidL Lipocalin |
| *Salmonella typhimurium* | STM3759;MarT | STM3758* | FidL Lipocalin |
| *Salmonella typhimurium* | STM0341 | STM0342* | half barrel |
| *Salmonella typhimurium* | STM0344 | STM0345* | Lipocalin |
| *Shigella flexneri* | SF3508 | SF3507* | Lipocalin |

\*Overlapping open reading frame

### VtrC defines a fast-evolving lipocalin-like sensor superfamily

The candidate co-component system gene cassettes include VtrA-like transcription factors and their adjacent periplasmic components. However, the adjacent periplasmic components tend to lack sequence similarity to known domains (42, 43). Given the ability of AlphaFold to confidently predict the fast-evolving ToxS structure, we generated models for each candidate VtrC-like co-component. For all but one case, the modeled structure adopted a lipocalin-like fold with eight meandering β-strands forming a VtrC-like barrel (Fig. 3*A*). Like the ToxS prediction (Fig. 2*D*), most of the models have high estimated confidence for residues in the core fold. However, the *Shigella* unknown protein SF3507, and the *Vibrio cholerae* TcpH models were less confident predictions. The final candidate, STM0342, was predicted as a four-stranded β-meander that corresponds to half of the barrel and may represent a deterioration of its sequence-related neighbor STM0345. Interestingly, STM0342 includes two Cys residues located too far from one another to form an intrachain disulfide, and the deterioration may form a homodimer suggested by an AlphaFold model (Fig. 3*B*).

**Figure 3.**
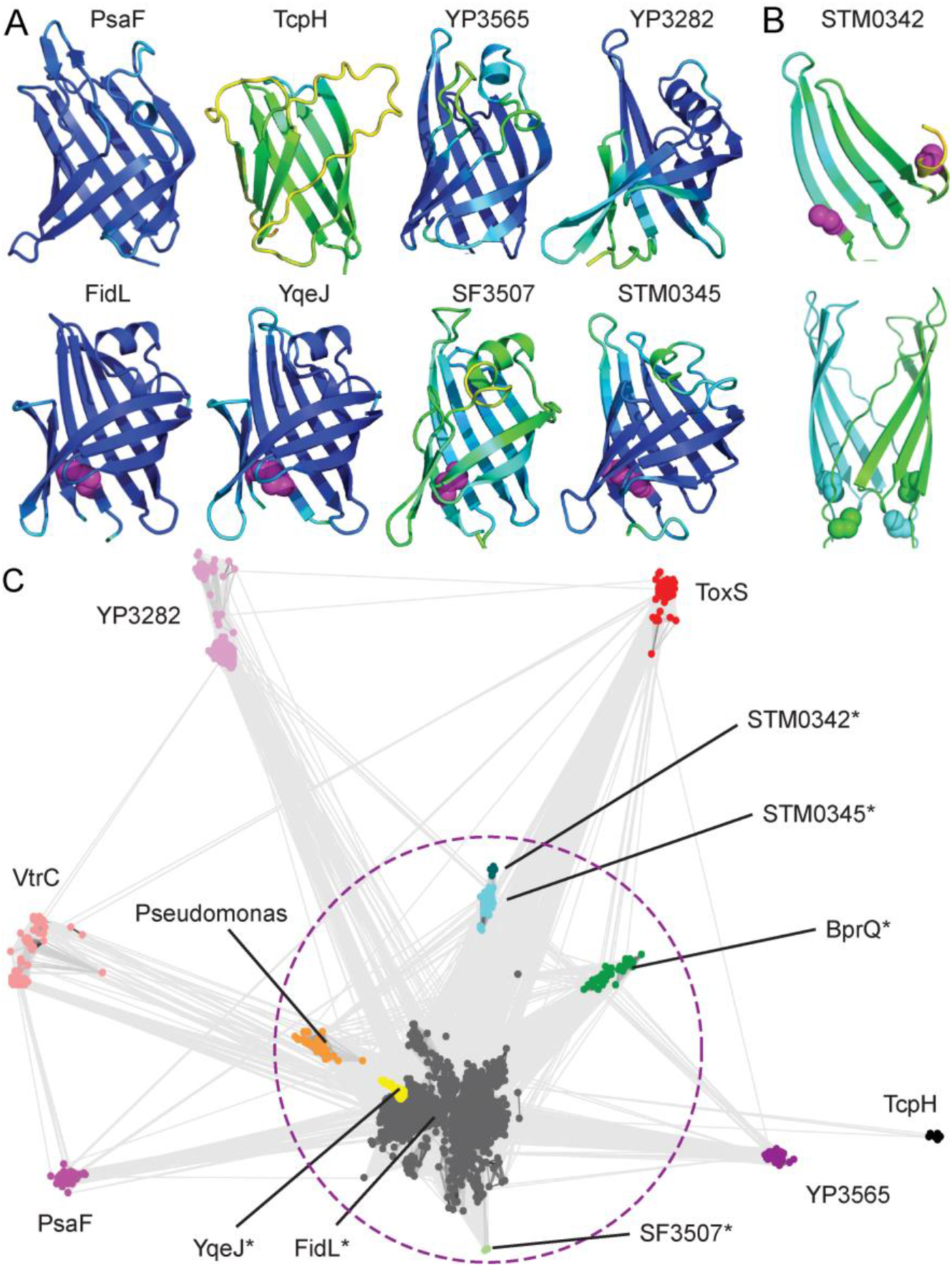
Lipocalin-like folds unite diverse VtrC-like members. **A)** Models for VtrC-like representatives are colored in rainbow from blue (high confidence) to red (low confidence), with Cys disulfides in magenta sphere. **B)** STM0342 monomer (above, depicted as in A) and dimer model (below, cyan and green) with cys residues (sphere). **C)** VtrC-like sequences (nodes, colored according to family and labeled) are clustered with CLANS in two dimensions. Connecting lines denote similarity between nodes (<0.0001 BLAST E-value cutoff). Families marked with (*) can be linked by sequence using PSI-BLAST and are circled with a dotted magenta line.

The lipocalin-like structure predictions for each of the VtrC-like components supports the notion that the second periplasmic component from each gene cassette arose from a common ancestor, despite their sequence divergence. In support of this proposed homologous relationship, the lipocalin-like folds from the *E. coli, Shigella* and *Salmonella* structure models possess a conserved disulfide formed between cysteine residues from the adjacent N-terminal and C-terminal strands of the barrel (Fig. 3*A*, magenta spheres), and the sequence relationship between *Salmonella* Spi-3 (*MarT/FidL*) and *E. coli* ETT2 (*YqeI/YqeJ*) has been noted (44). The presence of these conserved disulfides suggests some of the newly identified VtrC-like superfamily members might retain enough of a sequence signal to be recognized. To understand the sequence relationships between the identified families, we collected homologs for VtrC-like representatives and used these to build sequence profiles for each family. Profile comparisons using HHsearch (SI Appendix, Table S3) highlight the relationships between family members, where the disulfide-containing proteins FidL, YqeJ, STM0342, STM0345, and SF3507 are confidently related by sequence. The remaining families have diverged beyond confident sequence recognition using the most sensitive sequence comparison methods (HHsearch, SI Appendix, Table S3).

To visualize the relationships between all the VtrC-like families, collected sequence homologs were clustered (Fig. 3*C*). The disulfide-containing FidL-like sequences (Fig. 3*C*, gray nodes) overlap with the YqeJ-like sequences (Fig. 3*C*, yellow nodes) and are difficult to distinguish. Alternately, the clusters for the other confidently-related disulfide-containing families (i.e.SF3507, STM0342 and STM0345, Fig. 3*C*, enclosed by a dashed circle) are more separated. These diverging sequence-related clusters provide further support for the fast evolution of the superfamily. The cluster sizes (and distances from large clusters) of the various superfamily members tend to mimic the confidence values associated with the AlphaFold models, with TcpH (a singleton) being the least confident prediction. Thus, the presence of such sparsely populated and fast evolving families among the VtrC-like periplasmic components lacking the conserved disulfide has likely prevented their identification by traditional sequence-based methods.

Two additional groups of collected homologs are diverging from the FidL cluster, including BprQ-like sequences and sequences from various *Pseudomonas* species (Fig. 3*C*, green nodes and orange nodes, respectively). The BprQ cluster includes a T3SS3 regulator from the Melioidosis pathogen *Burkholderia pseudomallei*, which has recently been shown to be an enteric bacterium(45). The BprQ sequence adopts a predicted lipocalin-like fold with the preserved FidL-like disulfide. The *BprQ* gene is adjacent to *BprP*, which encodes a transmembrane anchored transcription factor. BprP/BprQ regulates the machinery and secretion components of the *Burkholderia pseudomallei* T3SS3(46, 47). BprP retains the same domain organization as other VtrA superfamily transcription factors, having an N-terminal HTH, followed by a TMH and a periplasmic region with the same predicted fold as the other superfamily members. The second detached cluster includes sequences from various other *Pseudomonas* strains, although the *Pseudomonas aeruginosa* PAO1 representative genome (SI Appendix, Table S2) lacks candidate co-component systems. One of the VtrC-like sequences from *Pseudomonas fluorescens* strain C1 overlaps with a VtrA-like transcription factor, highlighting the preserved operon organization in this species.

### Functional implications for the VtrA/VtrC-like co-component gene cassettes

The VtrA/VtrC periplasmic bile receptor forms an obligate heterodimer, and many of the identified candidate co-components are known to function together(13). Because protein-protein complex structure predictions can now be generated with increasing accuracy for larger families(31, 48, 49), we reasoned the sequence information from the larger FidL cluster (Fig. 3*B*) would be adequate for predicting its interaction with GrvA. Indeed, the structure model of the GrvA transcription factor in complex with FidL retains a similar interaction surface as in the VtrA/VtrC experimental structure, with top predicted residue-residue contacts lining the periplasmic domain interaction surface (Fig. 4*A*, yellow lines). The same β-barrel extension of a split observed in VtrC by the VtrA periplasmic β-sheet (Fig. 2*A*) is present in the FidL/GrvA complex (Fig. 4*A*). The TMH and relative position of the GrvA HTH are less confidently placed based on the predicted aligned error, and the per-residue confidence metric reflects lower accuracy in the TMH of both components as well as in the GrvA connecting helix and longer HTH domain loops (SI Appendix, Fig. S1). Protein-protein complex prediction for many of the other candidate co-components suggest similar interactions, although with generally lower confidence. In support of their heterodimeric interaction, the periplasmic domains from the co-component signal transduction systems PsaE/PsaF from *Yersinia* and YqeI/YqeJ from *E*.*coli* form dimeric complexes when expressed together (SI Appendix, Methods).

**Figure 4.**
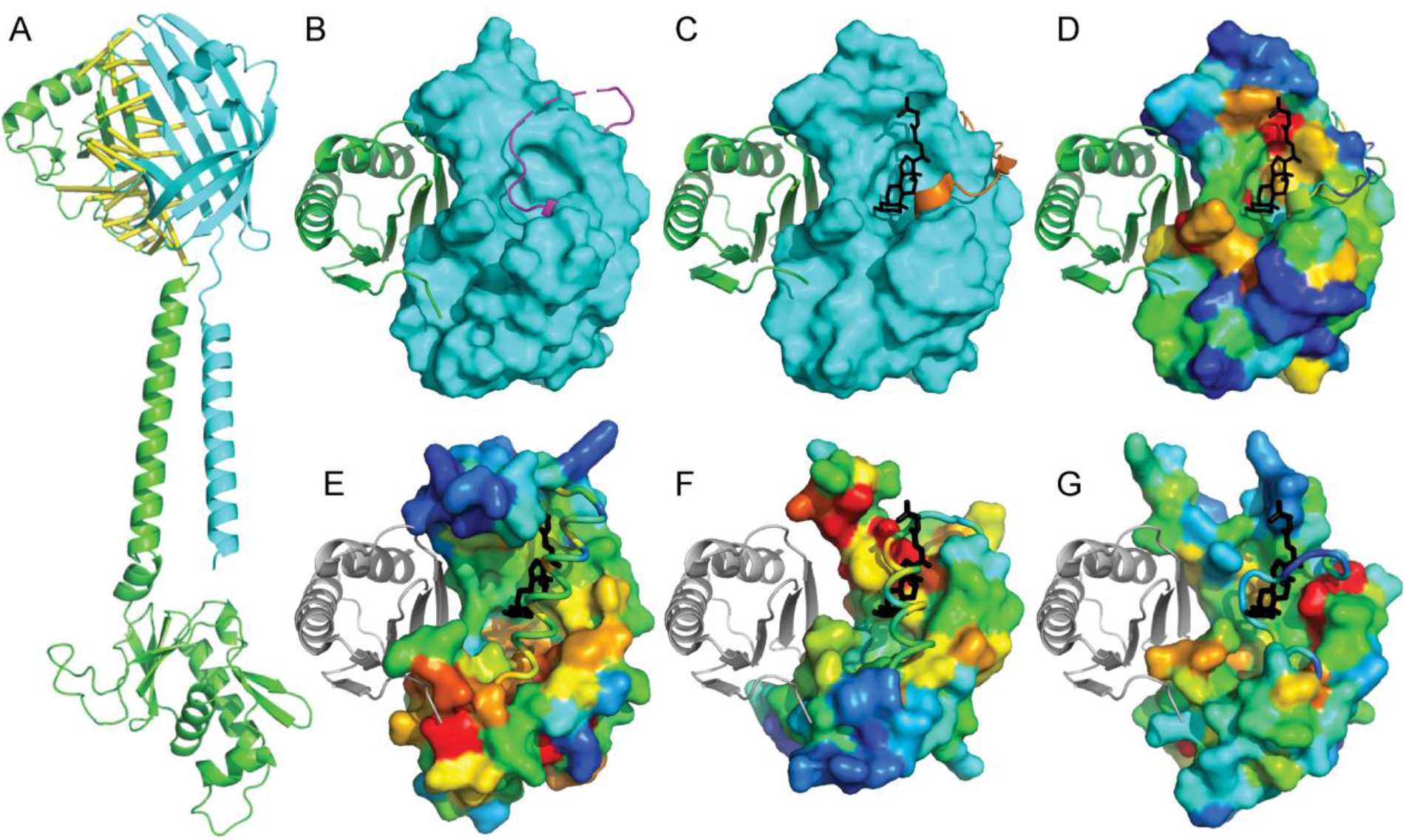
Functional implications of VtrA/VtrC superfamily fold prediction. **A)** GrvA (green cartoon) complex model with FidL (cyan cartoon). Confidently predicted residue-residue contacts are connected by yellow bars (residue-residue distance in structure model <= 8 Å and predicted aligned error <= 4 Å). **B)** VtrA (green cartoon) experimental structure bound to apo VtrC (cyan surface), with a mobile loop (orange) covering the lipid binding site. **C)** VtrA/VtrC experimental structure bound to TDC (black stick) displaces the mobile loop (magenta). **D)** VtrC surface is colored in rainbow by family-based residue conservation from low (blue) to high (red). **E)** ToxS **F)** YP3282 and **G)** BprQ structure models (rainbow conservation) are superimposed with VtrC to highlight relative positions of VtrA (white cartoon) and TDC (black stick). The corresponding mobile loops (cartoon tubes) cover the potential lipid binding sites as in the apo VtrA/VtrC structure.

The VtrA/VtrC periplasmic structure forms an obligate heterodimer with a mobile loop covering the lipid binding site in the absence of bile acid (Fig. 4*B*). Upon binding the bile acid TDC, the mobile loop opens to adopt an alternate conformation (Fig. 4*C*). Like other lipocalin folds(13), VtrC binds TDC on one side of the barrel center. This preserved binding mode, together with the lipocalin-like fold predictions of the other VtrC-like components, suggest they might also function as lipid sensors. Such functional sites are often conserved, and the resides that contribute to the binding site can be identified using multiple sequence alignments of family members. As a proof of this concept, conserved residue positions from VtrC contribute to the TDC binding site, and the VtrA interaction surface is also relatively conserved (Fig. 4*D*).

To gauge if the newly identified periplasmic components might bind lipid, we defined per-residue conservations for several of the VtrC-like clustered sequence groups and mapped them to the structure models to highlight potential functional surfaces. Superposition of the ToxS structure model with the experimental VtrC model bound to TDC (Fig. 4*E*) highlights a loop in ToxS that covers the potential lipid binding site. The loop corresponds to the mobile loop in VtrC, and it positions a helix in the corresponding lipid site. The covered lipid binding site is generally more conserved in ToxS than the surrounding surface, with the most conserved region surrounding the interior part of the cleft located deeper in the barrel. Portions of a potential ToxR binding site are represented by the superimposed VtrA periplasmic domain (Fig. 4*E*, white cartoon) and are particularly conserved at the surface interacting with the VtrA N-terminal loop. This conserved surface in ToxS might help position the ToxR TMH (connected to the loop), which could ultimately control the position of the intracellular DNA-binding domain.

Conservations in the YP3282 structure model (Fig. 4*F*) surround the exterior portion of the potential lipid binding cleft, which is also covered by a helix in the mobile loop. As opposed to the conservations in ToxS, the most conserved portion of the potential YP3282 transcription factor periplasmic domain surface lies on the sheet opposite to the N-terminal loop. Conservations in the BprQ structure model (Fig. 4*G*) also mark a potential lipid binding site that is covered by the mobile loop. However, the loop in BprQ is shorter and lacks a helix. Like the surface conservations in ToxS, the presumed BprP periplasmic domain binding site is most conserved near the N-terminal loop. Thus, while the various family members generally preserve conservations in the potential lipid binding site and the co-component interaction surface, these conservations differ between divergent members of the larger superfamily.

### Relationship between the VtrA-like co-components and CadC

The membrane embedded one-component transcription factor CadC includes an N-terminal HTH, followed by a TMH and a C-terminal periplasmic sensor (Fig. 1*B*). This domain organization is retained in the DNA-binding chain of the co-component systems (Fig. 1*C*). These similarities could suggest that the CadC and VtrA-like periplasmic domains arose by divergence from a common ancestor. The CadC one component periplasmic sensor includes two domains. The first adopts an αβ-sandwich, and the second adopts an α-helical domain of TPR repeats (Fig. 1*B*). The structure of the CadC αβ-sandwich (Fig. 5*A*) includes an N-terminal α/β/α unit with a Rossmann-like crossover, followed by a 3-stranded β-meander and a C-terminal α-helix. If the β-strand and loop from the Rossmann-like crossover represents an insert, the CadC periplasmic sandwich retains the same core set of secondary structure elements as in the VtrA co-component periplasmic domain (Fig. 5*B*), with an N-terminal β-strand extension in VtrA replacing the Rossmann-like insertion strand from CadC. Superposition of the two domains results in structure similarity (Dali Z-score 5.6) over the βαβ(3)α core that is above the suggested cutoff for significance (Z-score 2) and is higher than the corresponding structure similarity between the related ToxR and VtrA periplasmic domains (Z-score 4.1). However, these scores reside within the “gray area” of homology, and many existing structures exhibit higher similarities to the CadC fold, as VtrA is ranked 34 a search against a non-redundant structure database (PDB25)(50, 51).

**Figure 5.**
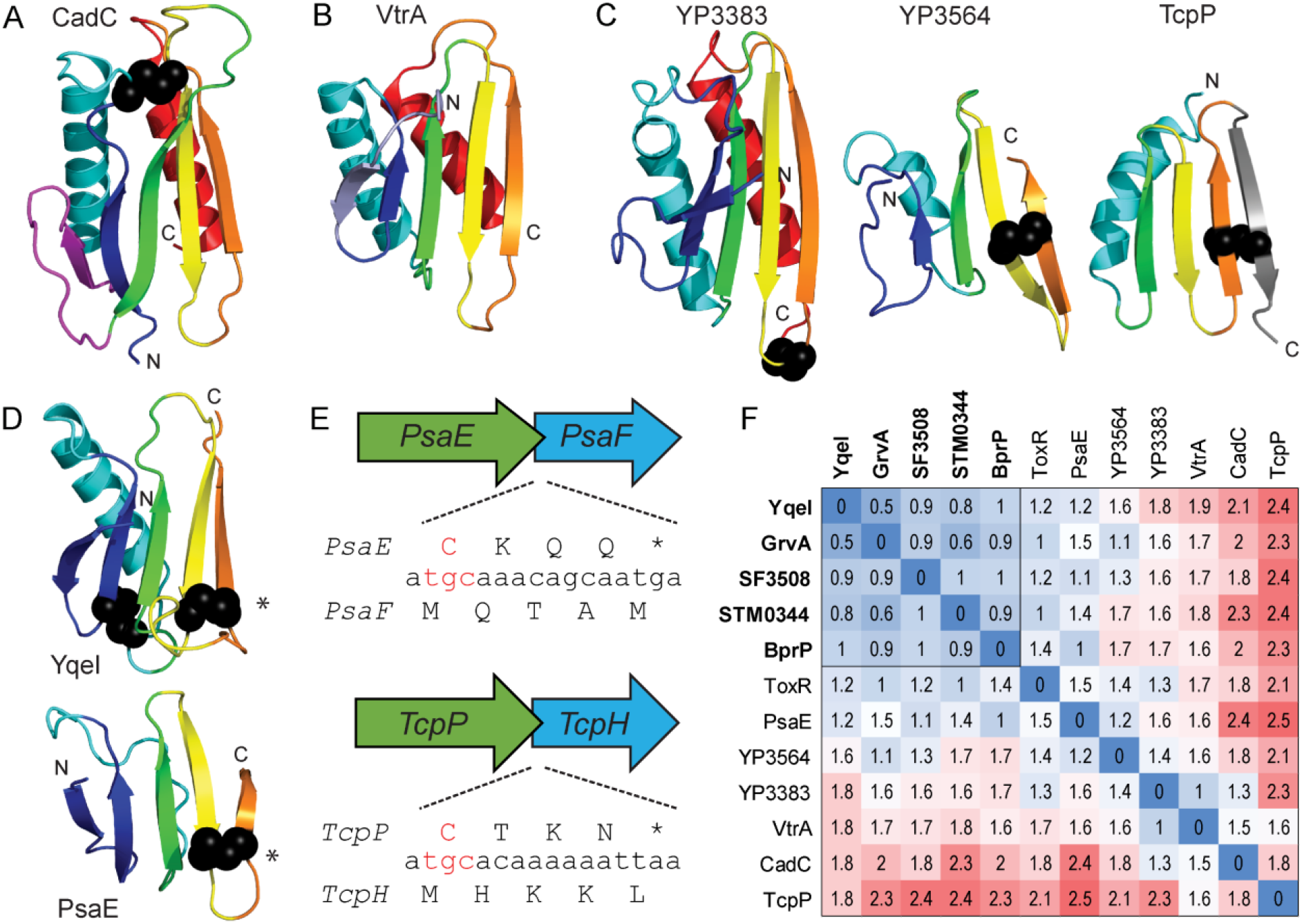
VtrA periplasmic domain evolution. Periplasmic domains SSEs are in rainbow cartoon from the N-terminus (blue) to the C-terminus (red), with disulfides in black sphere. **A)** CadC cadaverine synthesis transcription sandwich domain has an insertion (magenta) with respect to **B)** the VtrA periplasmic domain, with the CadC insertion replaced by an N-terminal extension (slate). **C)** Candidate co-component VtrA-like models are colored by the core SSEs in CadC and VtrA. **D)** A conserved disulfide (*) links YqeI and TcpP, which includes a C-terminal extension (gray). **E)** Representative operon ORFs (labeled) with conserved C-terminal Cys codon (red) overlapping with start codon from input component. **F)** Distance matrix of all-against-all structure comparisons colored by distance in red (distant) - white - bue (identical) color scale. Components that interact with confident sequence-related VtrC-like domains are bold and their distances are boxed.

The lack of sequence similarity combined with marginal structure similarity between the CadC and VtrA periplasmic domains precludes sequence-based clustering or traditional phylogenetic analysis to evaluate evolutionary relationships. However, the models for the VtrA-like periplasmic domains allow quantitative comparison of the structures as well as definition of the conserved SSEs that contribute to the fold. The experimental VtrA and ToxR periplasmic domain structures include a common set of SSEs (β(2)αβ(3)α) that can be compared to the rest of the VtrA-like components (Fig. 5*C and D*). The *Yersinia* YP3564 model is the only one with the C-terminal helix found in VtrA and ToxR, while its distantly related paralog (YP3383) model lacks the helix and shows a deteriorating N-terminal strand. The models for several of the VtrA-like components (YqeI, GrvA, SF3508, STM0344, and BprP) retain a similar overall fold without the C-terminal helix (YqeI in Fig. 5*D*). The N-terminal helix has also deteriorated into a connecting loop in the PsaE model (Fig. 5*D*). Finally, the model for TcpH lacks both N-terminal β-strands and gains a C-terminal β-strand. The interaction mode for VtrA/VtrC co-component system uses the C-terminal strand to extend a sheet in the opened VtrC barrel. The lack of a C-terminal helix would better expose the C-terminal edge strand for such an interaction (as in Fig. 2*A* and 4*A*), and the deterioration of SSEs on the opposite side is consistent with their distance from the functional interaction surface. However, the C-terminal extension of the sheet in TcpP would not be compatible with the VtrA/VtrC-like interaction.

Notably, except for VtrA, the co-component transcription factor periplasmic domains contain conserved disulfides. Each of these includes a Cys residue near the C-terminus of the protein sequence (Fig. 5*B* and Fig. 5*D*). With a few exceptions (ToxR/ToxS, YqeI/YqeJ and GrvA/FidL), the transcription factor periplasmic domain C-terminus overlaps with the N-terminus of the neighboring co-component gene, which is encoded by a frame shift (Fig. 5*E* and SI Appendix, Fig. S3). The starting Met (atg codon) from the downstream gene tends to overlap with the Cys (tg(t/c) codon) from the disulfide. In this case the starting Met from the second gene is shifted by one nucleotide upstream with respect to the first gene. The *TcpP/TcpH, PsaE/PsaF, YP3564/YP3565, STM0341/STM0342*, and *STM0344/STM0345* gene overlaps observe this tendency. Alternately, SF3508/SF3507 and YP3283/YP3282, which are encoded in the complementary strands, include the Cys residues in their overlap. Finally, the MarT/FidL overlap includes a His residue (cat codon) shifted one nucleotide upstream with respect to the MarT overlapping Met (atg codon).

A heatmap of distances calculated from pair-wise comparisons of the VtrA-like periplasmic domain structures highlights their relationships (Fig. 5*F*). A subset of VtrA-like periplasmic domains (YqeI, GrvA, stm0344, SF3808, and BprP) that interact with the sequence related VtrC-like components retain relatively lower distances. These domains include a conserved disulfide bond between the last two β-strands (Fig. 5*D*, bold labels). This disulfide bond retention extends the confident assignment of co-component homologs to include the PsaE/PsaF cassette, and by transitivity to the VtrA/VtrC cassette, given the detected sequence relationship between their interacting partners VtrC and PsaF (SI Appendix, Table S1). The more divergent VtrA-like components (Fig. 5*F*) interact with the more divergent VtrC-like components (Fig. 3*B*). The CadC subdomain distances to the rest of the co-component periplasmic domains tend to be less than those of TcpP, and its average similarity to the rest of the structures is higher (Z-score 3.16 for CadC and 1.55 for TcpP). The closest co-component structure to CadC is in the gray zone of homology (VP3383: Z-score 6.7). The distance heatmap resembles a heatmap generated using similarity scores from another structure superposition program (SI Appendix, Figure S2). Although tempting to speculate the co-component signal transduction systems replaced the loss of the CadC helical domain with a gained VtrC-like heterodimer, their structure dissimilarity prevents any confident assignment of homology. This questionable relationship extends to the TcpP component, especially given the tendency of domain recombination among bacterial signaling proteins(6, 7) and the presumed interaction mode of the co-components.

## Discussion

While similarity between the VtrA, ToxR and other CadC-like HTH domains has long been known (38, 52), sequence divergence in the periplasmic regions has hampered classification. Similarly, while the functional association between the transcription factors and their neighboring gene products is either known or implied (in some cases, the open reading frame is not even annotated), fast sequence evolution of the co-activators has precluded their assignment to known structures or functions. The structure predictions for VtrA/VtrC-like superfamily suggest the co-component systems arose from a common two-gene cassette ancestor that has distributed across various enteric bacteria. One of the genes encodes a membrane-tethered transcription factor output domain, and the other encodes a lipid binding-like input co-sensor. The VtrA/VtrC co-components function as an obligate heterodimer, where the incomplete barrel for VtrC is unstable. Similarly, the ToxR periplasmic domain is regulated by its stability, which is influenced both by interaction with ToxS and by formation of a disulfide (18, 22, 24). Thus, ToxS is thought to activate ToxR by protecting it from degradation(53). Similar chaperone-like mechanisms are proposed for the TcpP/TcpH and PsaE/PsaF co-components (54, 55). While degradation might represent one mode of regulation for these co-component systems, their evolutionary relationship to the VtrA/VtrC bile acid sensor suggests they might form heterodimers and respond to a lipid-like environmental cues.

Enteric bacteria exhibit a dual relationship with bile for survival in the human gastrointestinal tract. On the one hand, both commensal and pathogenic microorganisms must contend with the antimicrobial actions of bile. On the other hand, pathogenic bacteria can use bile as an environmental cue to generate virulence factors(56). The *Vibrio cholera* master virulence regulator ToxR/ToxS is responsive to bile, yet the detailed mechanism of transcription activation by bile remains elusive (16, 18, 20, 23, 24). A complex regulatory cascade, known as the ToxR/ToxS regulon, ultimately produces the virulence toxin CtxAB and the toxin coregulated pilus TcpA. This complexity confounds understanding the contribution of bile to regulation. For example, the downstream ToxR/ToxS responsive ToxT regulator represses the regulon in response to arachidonic, linoleic, and oleic acid components of bile, and a structure of ToxT bound to palmitoleic acid supports these observations (57, 58). The ToxR/ToxS regulon is enhanced by other bile constituents. The bile acid taurocholate activates the ToxR/ToxS regulon by promoting TcpP dimerization (59) and ToxR/ToxS itself responds to deoxycholate by increased ToxR-ToxR interactions (23). These data are consistent with TcpH and ToxS sensing bile and activating transcription through their TcpP and ToxR co-component DNA binding domains.

The relationship between the remaining co-component systems and lipid binding are less clear. The *S*. Typhimurium MarT/FidL co-components belong to the Spi-3 pathogenicity island and regulate expression of their neighboring *MisL* gene. The *MisL* gene product encodes an autotransporter that functions as an intestinal colonization factor (38-40). While this function would be consistent with FidL sensing an environmental cue from the intestine, the mechanism of transcriptional activation by MarT remains to be determined. Interestingly, the *MarT* and *MisL* genes have acquired stop codons and become pseudogenized in the *S*. Typhi serovar. The loss of MarT/MisL function in the *S*. Typhi background helped improved virulence in a restricted host cell range (60, 61). Genome comparisons have revealed a distant relationship between the Spi-3 pathogenicity island and the ETT2 type III secretion system in *E. coli* and *Shigella* (44). This relationship identifies *E*.*coil* K12 and Saki YqeI/YqeJ as orthologs of *Salmonella* MarT/FidL. The corresponding gene region is missing in the incomplete ETT2 gene cluster from *Shigella*, although a paralogous uncharacterized co-component cassette exists elsewhere in the *S. flexneri* genome (SF3508/SF3507). The *E*.*coil* K12 YqeI/YqeJ cassette also appears to be a remnant of the complete ETT2 pathogenicity island present in pathogenic strains like *E. coli* O157:H7 Saki. Pathogenic *E. coli* O157:H7 Saki includes a paralogous YqeI/YqeJ gene cassette (*GrvA/FidL*) that activates the locus of enterocyte effacement (LEE)-dependent adherence (41), and many of the LEE adherence genes are differentially regulated by the presence of bile acids (62).

The similarity in folds between the CadC N-terminal periplasmic domain and the periplasmic domains from the co-component transcription factors points towards their derivation from a common membrane-tethered transcription factor. The Cad system, which is widespread in enteric bacteria, helps maintain a neutral cytoplasmic pH in the acidic environment of the stomach (10, 12). The CadC transcription factor is activated in part by acid conditions, which reduce the N-terminal periplasmic domain disulfide (63). Some of the co-component transcription factors such as ToxR are also known to respond to pH. However, ToxR is activated in alkaline conditions and requires oxidized disulfides (23).

Although the positions of disulfides in the periplasmic domains of these transcription factors are not preserved (fig. 5*E*), they might point to a common function in pH sensing. In fact, a disulfide stabilizes the unusual C-terminal strand topology of TcpP. Potentially, low pH might reduce the disulfide and release the strand to allow interaction with TcpH. In fact, the solution structure of oxidized ToxR revealed similar structure instability in the C-terminus where the periplasmic domain presumably interacts with its ToxS partner. Such structure plasticity might help explain the fast evolution of the periplasmic domain, as the unstructured protein regions can evolve independent of their primary sequence if they maintain the disulfide. Apart from VtrA, which lacks a disulfide, the co-component transcription factors maintain a disulfide Cys residue near the C-terminus (Fig. 5*E*). This position tends to correspond with the sequence overlap from the neighboring gene (SI Appendix, Fig. S3), providing a mechanism for its preservation.

Overall, our studies support a model for a new signaling paradigm used by enteric bacteria to sense extracellular cues in the periplasm. The co-component signal transduction system includes an input periplasmic sensor that is tightly associated with its transcription factor, as exhibited by their operon organization, co-evolution and obligate periplasmic domain interaction. This co-component system provides an additional, albeit specialized, mechanism for transmitting an extracellular signal sensed by a periplasmic input through the membrane to a transcription factor output domain, adding to the canonical transmembrane one-component and two-component signaling systems (Fig. 1). Understanding the mechanism of signal transduction at the membrane and the ligands or conditions that activate these co-components will provide seminal information on how bacteria respond to their environment.

## Methods

### Extending the VtrA/VtrC co-component superfamily

Sequence corresponding to *Vibrio parahaemolyticus* ToxR (BAC59083.1) and VtrA (BAC62675.1) were used as queries with PSI-BLAST(64) (5 iterations, E-value cutoff 0.001) to search against a protein sequence database comprised of bacterial reference genomes. The database was constructed using makeblastdb with protein sequences from RefSeq reference genomes defined by NCBI (15 genomes) together with protein sequences from the *Vibrio parahaemoliticus* RIMD, *Vibrio cholera* MS6, and *Yersinia pestis* A1122 genomes with gene accessions in NCBI (SI Appendix, Table S2). Identified VtrA-related hits were filtered for a predicted TMH using Phobius (65). The genome neighborhoods of candidate TMH-containing VtrA-like sequences (34 genes) were inspected for a tandem operon of the VtrA-like sequence and a downstream potential VtrC-like sequence with a predicted N-terminal TMH or signal peptide (SP). To account for potential overlapping ORFs (as is the case for VtrA/VtrC), the upstream in-frame region was translated for cases where the adjacent gene is missing a predicted N-terminal TMH or SP.

### VtrC-like Sequence Clustering and Family-Level Conservations

For clustering VtrC-like sequences, close sequence homologs of each of the VtrC-like family members with a predicted lipid binding domain were used as queries to search the RefSeq Select database using the NCBI server (default values, search until convergence, or relaxed E-value cutoff 0.02 for small families). All sequences were clustered with CLANS(66) using BLAST scores (E-value cutoff 0.01). Identified sequences were submitted to the MAFFT server(67) to generate multiple sequence alignments with the default strategy. Residue conservations were mapped to the B-factors of the VtrC structure (5kew, chain D) or the structure models of ToxS and FidL using Al2Co(68) for visualization in PyMOL with a rainbow color scale from blue (variable) to red (conserved).

### Single and Complex Structure Prediction

Candidate VtrC-like sequences found in tandem with VtrA-like transmembrane transcription factors were submitted to AlphaFold2(32) structure prediction using the ColabFold(35)), which replaces the homology detection of AlphaFold2 with MMseqs2(69), or with a local adaptation of AlphaFold described below.

Starting from candidate co-component protein pairs, we searched for the homologs of each protein encoded by nucleotide sequences in the European Nucleotide Archive database and the Integrated Microbial Genomes and Microbiomes database of the Joint Genome Institute. Six rounds of iterative HMMER (70) search with e-value cut-offs of 10-12, 10-12, 10-12, 10-12, 10-6, 10-3, respectively were used. Homologs found in each round of sequence search were used to construct the sequence profile for each protein using HMMER hmmbuild, which was used to identify more homologs in the next round. We filtered the homologs found in the last round of database search by their coverage (> 60 %) over the query sequence and recorded their loci on the nucleotide sequences.

For each protein pair, their homologs that are encoded next to each other in the nucleotide sequences were extracted. The sequences of these protein pairs were concatenated, and the multiple sequence alignment (MSA) was derived from the pairwise sequence alignments made by HMMER. The MSA was then filtered by sequence identity (maximal identify for remaining sequences <= 90 %) and gap ratio in each sequence (maximal gap ratio <= 50%), and the resulting non-redundant MSA was used as input for complex structure modelling.

We deployed AlphaFold (32), a method designed to model protein monomers, to model protein complexes using two modifications: (1) we provided the concatenated alignments of protein pairs (described above) to AlphaFold instead of sequence alignments for single proteins, (2) we changed the positional encoding used in ALPHAFOLD to represent a chain break between two proteins. The relative positional encoding is calculated based on residue numbers and it is the only input that encodes chain connectivity in AlphaFold. We added 200 to the residue numbers of the second protein in each concatenated alignment to let the AlphaFold network know that there is no chain connection between the two proteins. This simple modification has enabled us to model a large set of Eukaryotic protein complexes [34762488], and it is expected to produce high-quality model for candidate one-component signal transduction systems with enough homologs (∼100).

### VtrA-like periplasmic domain heatmap

The experimental structures corresponding to the VtrA periplasmic domain (5kewA), the ToxR periplasmic domain (6uueA), and the CadC N-terminal periplasmic domain (3lyaA, residues 194-316) were compared to the models for GrvA (189-270), YqeI (186-269), STM0344 (170-247), SF3808 (170-254), BprP (256-326), PsaE (160-214), YP3564 (162-219), YP3383 (235-348), and TcpH (169-221). All-against-all structures were compared using DaliLte (71), Pairwise (Z_AB_) and self (Z_AA_, Z_BB_) Z-scores were transformed into distances using the following equation: -ln(Z_AB_/(minimum of Z_AA_,Z_BB_)).

## Acknowledgements

We thank the Orth lab members for discussions and editing. This work was funded by the Welch Foundation grant I-1561 (K.O.), Once Upon a Time…Foundation (K.O.), and National Institutes of Health R01 GM115188 (K.O.). K.O. is a W.W. Caruth, Jr. Biomedical Scholar with an Earl A. Forsythe Chair in Biomedical Science. Q. C. is a Southwestern Medical Foundation Scholar.

## References

1. M. A. Matilla, A. Ortega, T. Krell, The role of solute binding proteins in signal transduction. Comput Struct Biotechnol J 19, 1786–1805 (2021).

2. C. Park, G. L. Hazelbauer, Mutations specifically affecting ligand interaction of the Trg chemosensory transducer. J Bacteriol 167, 101–109 (1986).

3. U. S. Cho et al., Metal bridges between the PhoQ sensor domain and the membrane regulate transmembrane signaling. J Mol Biol 356, 1193–1206 (2006).

4. E. A. Groisman, The pleiotropic two-component regulatory system PhoP-PhoQ. J Bacteriol 183, 1835–1842 (2001).

5. J. A. Hoch, Two-component and phosphorelay signal transduction. Curr Opin Microbiol 3, 165– 170 (2000).

6. L. E. Ulrich, E. V. Koonin, I. B. Zhulin, One-component systems dominate signal transduction in prokaryotes. Trends Microbiol 13, 52–56 (2005).

7. K. Wuichet, B. J. Cantwell, I. B. Zhulin, Evolution and phyletic distribution of two-component signal transduction systems. Curr Opin Microbiol 13, 219–225 (2010).

8. G. Rivera-Cancel, K. Orth, Biochemical basis for activation of virulence genes by bile salts in Vibrio parahaemolyticus. Gut Microbes 8, 366–373 (2017).

9. S. Buchner, A. Schlundt, J. Lassak, M. Sattler, K. Jung, Structural and Functional Analysis of the Signal-Transducing Linker in the pH-Responsive One-Component System CadC of Escherichia coli. J Mol Biol 427, 2548–2561 (2015).

10. C. L. Dell, M. N. Neely, E. R. Olson, Altered pH and lysine signalling mutants of cadC, a gene encoding a membrane-bound transcriptional activator of the Escherichia coli cadBA operon. Mol Microbiol 14, 7–16 (1994).

11. A. Eichinger, I. Haneburger, C. Koller, K. Jung, A. Skerra, Crystal structure of the sensory domain of Escherichia coli CadC, a member of the ToxR-like protein family. Protein Sci 20, 656–669 (2011).

12. A. G. Torres, The cad locus of Enterobacteriaceae: more than just lysine decarboxylation. Anaerobe 15, 1–6 (2009).

13. P. Li et al., Bile salt receptor complex activates a pathogenic type III secretion system. Elife 5 (2016).

14. A. Ortega, I. B. Zhulin, T. Krell, Sensory Repertoire of Bacterial Chemoreceptors. Microbiol Mol Biol Rev 81 (2017).

15. B. M. Childers, K. E. Klose, Regulation of virulence in Vibrio cholerae: the ToxR regulon. Future Microbiol 2, 335–344 (2007).

16. V. L. Miller, R. K. Taylor, J. J. Mekalanos, Cholera toxin transcriptional activator toxR is a transmembrane DNA binding protein. Cell 48, 271–279 (1987).

17. C. W. Ronson, B. T. Nixon, F. M. Ausubel, Conserved domains in bacterial regulatory proteins that respond to environmental stimuli. Cell 49, 579–581 (1987).

18. N. Gubensak et al., The periplasmic domains of Vibriocholerae ToxR and ToxS are forming a strong heterodimeric complex independent on the redox state of ToxR cysteines. Mol Microbiol 115, 1277–1291 (2021).

19. C. R. Midgett, R. A. Swindell, M. Pellegrini, F. Jon Kull, A disulfide constrains the ToxR periplasmic domain structure, altering its interactions with ToxS and bile-salts. Sci Rep 10, 9002 (2020).

20. V. L. Miller, V. J. DiRita, J. J. Mekalanos, Identification of toxS, a regulatory gene whose product enhances toxR-mediated activation of the cholera toxin promoter. J Bacteriol 171, 1288–1293 (1989).

21. V. J. DiRita, J. J. Mekalanos, Periplasmic interaction between two membrane regulatory proteins, ToxR and ToxS, results in signal transduction and transcriptional activation. Cell 64, 29–37 (1991).

22. S. Almagro-Moreno, M. Z. Root, R. K. Taylor, Role of ToxS in the proteolytic cascade of virulence regulator ToxR in Vibrio cholerae. Mol Microbiol 98, 963–976 (2015).

23. M. Lembke et al., Host stimuli and operator binding sites controlling protein interactions between virulence master regulator ToxR and ToxS in Vibrio cholerae. Mol Microbiol 114, 262–278 (2020).

24. C. R. Midgett et al., Bile salts and alkaline pH reciprocally modulate the interaction between the periplasmic domains of Vibrio cholerae ToxR and ToxS. Mol Microbiol 105, 258–272 (2017).

25. D. T. Hung, J. J. Mekalanos, Bile acids induce cholera toxin expression in Vibrio cholerae in a ToxT-independent manner. Proc Natl Acad Sci U S A 102, 3028–3033 (2005).

26. T. Kodama et al., Two regulators of Vibrio parahaemolyticus play important roles in enterotoxicity by controlling the expression of genes in the Vp-PAI region. PLoS One 5, e8678 (2010).

27. R. L. Dunbrack, Jr., Sequence comparison and protein structure prediction. Curr Opin Struct Biol 16, 374–384 (2006).

28. J. Moult, A decade of CASP: progress, bottlenecks and prognosis in protein structure prediction. Curr Opin Struct Biol 15, 285–289 (2005).

29. W. R. Pearson, M. L. Sierk, The limits of protein sequence comparison? Curr Opin Struct Biol 15, 254–260 (2005).

30. L. N. Kinch, R. D. Schaeffer, A. Kryshtafovych, N. V. Grishin, Target classification in the 14th round of the critical assessment of protein structure prediction (CASP14). Proteins 10.1002/prot.26202 (2021).

31. M. Baek et al., Accurate prediction of protein structures and interactions using a three-track neural network. Science 373, 871–876 (2021).

32. J. Jumper et al., Highly accurate protein structure prediction with AlphaFold. Nature 596, 583–589 (2021).

33. L. N. Kinch, J. Pei, A. Kryshtafovych, R. D. Schaeffer, N. V. Grishin, Topology evaluation of models for difficult targets in the 14th round of the critical assessment of protein structure prediction. Proteins 10.1002/prot.26172 (2021).

34. K. Tunyasuvunakool et al., Highly accurate protein structure prediction for the human proteome. Nature 596, 590–596 (2021).

35. M. Mirdita, S. Ovchinnikov, M. Steinegger, ColabFold - Making protein folding accessible to all. bioRxiv 10.1101/2021.08.15.456425, 2021.2008.2015.456425 (2021).

36. Y. Yang, R. R. Isberg, Transcriptional regulation of the Yersinia pseudotuberculosis pH6 antigen adhesin by two envelope-associated components. Mol Microbiol 24, 499–510 (1997).

37. C. C. Hase, J. J. Mekalanos, TcpP protein is a positive regulator of virulence gene expression in Vibrio cholerae. Proc Natl Acad Sci U S A 95, 730–734 (1998).

38. A. B. Blanc-Potard, F. Solomon, J. Kayser, E. A. Groisman, The SPI-3 pathogenicity island of Salmonella enterica. J Bacteriol 181, 998–1004 (1999).

39. C. W. Dorsey, M. C. Laarakker, A. D. Humphries, E. H. Weening, A. J. Baumler, Salmonella enterica serotype Typhimurium MisL is an intestinal colonization factor that binds fibronectin. Mol Microbiol 57, 196–211 (2005).

40. C. Tukel et al., MarT activates expression of the MisL autotransporter protein of Salmonella enterica serotype Typhimurium. J Bacteriol 189, 3922–3926 (2007).

41. J. K. Morgan et al., Global Regulator of Virulence A (GrvA) Coordinates Expression of Discrete Pathogenic Mechanisms in Enterohemorrhagic Escherichia coli through Interactions with GadW-GadE. J Bacteriol 198, 394–409 (2016).

42. S. Lu et al., CDD/SPARCLE: the conserved domain database in 2020. Nucleic Acids Res 48, D265–D268 (2020).

43. J. Mistry et al., Pfam: The protein families database in 2021. Nucleic Acids Res 49, D412–D419 (2021).

44. C. P. Ren et al., The ETT2 gene cluster, encoding a second type III secretion system from Escherichia coli, is present in the majority of strains but has undergone widespread mutational attrition. J Bacteriol 186, 3547–3560 (2004).

45. A. Goodyear, H. Bielefeldt-Ohmann, H. Schweizer, S. Dow, Persistent gastric colonization with Burkholderia pseudomallei and dissemination from the gastrointestinal tract following mucosal inoculation of mice. PLoS One 7, e37324 (2012).

46. G. W. Sun et al., Identification of a regulatory cascade controlling Type III Secretion System 3 gene expression in Burkholderia pseudomallei. Mol Microbiol 76, 677–689 (2010).

47. N. R. Lazar Adler et al., Perturbation of the two-component signal transduction system, BprRS, results in attenuated virulence and motility defects in Burkholderia pseudomallei. BMC Genomics 17, 331 (2016).

48. I. R. Humphreys et al., Computed structures of core eukaryotic protein complexes. Science 374, eabm4805 (2021).

49. R. Evans et al., Protein complex prediction with AlphaFold-Multimer. bioRxiv 10.1101/2021.10.04.463034, 2021.2010.2004.463034 (2021).

50. L. Holm, DALI and the persistence of protein shape. Protein Sci 29, 128–140 (2020).

51. L. Holm, Using Dali for Protein Structure Comparison. Methods Mol Biol 2112, 29–42 (2020).

52. D. S. Merrell, A. Camilli, Regulation of vibrio cholerae genes required for acid tolerance by a member of the “ToxR-like” family of transcriptional regulators. J Bacteriol 182, 5342–5350 (2000).

53. M. Lembke et al., Proteolysis of ToxR is controlled by cysteine-thiol redox state and bile salts in Vibrio cholerae. Mol Microbiol 110, 796–810 (2018).

54. S. J. Morgan et al., Formation of an Intramolecular Periplasmic Disulfide Bond in TcpP Protects TcpP and TcpH from Degradation in Vibrio cholerae. J Bacteriol 198, 498–509 (2016).

55. J. D. Quinn, E. H. Weening, V. L. Miller, PsaF Is a Membrane-Localized pH Sensor That Regulates psaA Expression in Yersinia pestis. J Bacteriol 203, e0016521 (2021).

56. M. Begley, C. G. Gahan, C. Hill, The interaction between bacteria and bile. FEMS Microbiol Rev 29, 625–651 (2005).

57. A. Chatterjee, P. K. Dutta, R. Chowdhury, Effect of fatty acids and cholesterol present in bile on expression of virulence factors and motility of Vibrio cholerae. Infect Immun 75, 1946–1953 (2007).

58. M. J. Lowden et al., Structure of Vibrio cholerae ToxT reveals a mechanism for fatty acid regulation of virulence genes. Proc Natl Acad Sci U S A 107, 2860–2865 (2010).

59. M. Yang et al., Bile salt-induced intermolecular disulfide bond formation activates Vibrio cholerae virulence. Proc Natl Acad Sci U S A 110, 2348–2353 (2013).

60. P. Retamal, M. Castillo-Ruiz, N. A. Villagra, J. Morgado, G. C. Mora, Modified intracellular-associated phenotypes in a recombinant Salmonella Typhi expressing S. Typhimurium SPI-3 sequences. PLoS One 5, e9394 (2010).

61. A. P. Ortega et al., Lose to win: marT pseudogenization in Salmonella enterica serovar Typhi contributed to the surV-dependent survival to H2O2, and inside human macrophage-like cells. Infect Genet Evol 45, 111–121 (2016).

62. X. Yin et al., Differential gene expression and adherence of Escherichia coli O157:H7 in vitro and in ligated pig intestines. PLoS One 6, e17424 (2011).

63. L. Tetsch, C. Koller, A. Donhofer, K. Jung, Detection and function of an intramolecular disulfide bond in the pH-responsive CadC of Escherichia coli. BMC Microbiol 11, 74 (2011).

64. S. F. Altschul et al., Gapped BLAST and PSI-BLAST: a new generation of protein database search programs. Nucleic Acids Res 25, 3389–3402 (1997).

65. L. Kall, A. Krogh, E. L. Sonnhammer, Advantages of combined transmembrane topology and signal peptide prediction--the Phobius web server. Nucleic Acids Res 35, W429–432 (2007).

66. T. Frickey, A. Lupas, CLANS: a Java application for visualizing protein families based on pairwise similarity. Bioinformatics 20, 3702–3704 (2004).

67. K. Katoh, K. Misawa, K. Kuma, T. Miyata, MAFFT: a novel method for rapid multiple sequence alignment based on fast Fourier transform. Nucleic Acids Res 30, 3059–3066 (2002).

68. J. Pei, N. V. Grishin, AL2CO: calculation of positional conservation in a protein sequence alignment. Bioinformatics 17, 700–712 (2001).

69. M. Steinegger, J. Soding, MMseqs2 enables sensitive protein sequence searching for the analysis of massive data sets. Nat Biotechnol 35, 1026–1028 (2017).

70. L. S. Johnson, S. R. Eddy, E. Portugaly, Hidden Markov model speed heuristic and iterative HMM search procedure. BMC Bioinformatics 11, 431 (2010).

71. L. Holm, J. Park, DaliLite workbench for protein structure comparison. Bioinformatics 16, 566–567 (2000).

