## Supplementary material for "Co-Component Signal Transduction Systems: fast-evolving virulence regulation cassettes discovered in enteric bacteria": SI Appendix

**Supplementary Information for**

### Co-Component Receptors: Fast-evolving virulence regulating cassettes in enteric bacteria discovered with the VtrA/VtrC operon.

Lisa N. Kinch, Qian Cong, Jananee Jaishankar and Kim Orth*

Corresponding Author: Kim Orth Inaugural Article

#### Supplementary Information Text

**Protein Expression and Purification.** Given the predicted heterodimeric interactions of the co-component receptors, we propose that, like the VtrA/VtrC co-component receptor, when the periplasmic domains of co-component receptors are expressed together, they will form a dimeric complex. Consistent with this proposal, the periplasmic domains of the co-component receptors PsaE/PsaF from *Yersinia* and YqeI/YqeJ from *E.coli* form a complex upon expression. By tagging one of the two domains, we are able to co-purify the other component by affinity purification. In addition, gel filtration shows the complexes are stable and can be purified together.

#### Methods

##### Protein Cloning, Expression, and Purification

The periplasmic domains of YqeJ (aa 26-159) and YqeI (aa 180-269) were amplified from *E. coli* O157:H7 and cloned into the first and second MCS of modified pETDuet-1 vector (containing N-terminal hexahistidine tag followed by a DrICE cutting site (DEVDA) preceding the first MCS site), where YqeJ is N terminally hexahistidine-tagged. This construct was expressed in *E. coli* BL21 (DE3) cells. To induce protein expression, cultures were grown in 2xYT medium until OD600 reached 0.5-0.6 and induced with 0.4 mM isopropyl thiogalactopyranoside (IPTG) overnight at 25℃. Similarly, the periplasmic domains of PsaF (aa 22-162) and PsaE (aa 158-214) were amplified from *Yersinia pseudotuberculosis* YPIII strain and cloned into the first and second MCS of modified pETDuet-1 vector, where PsaF is N terminally hexahistidine-tagged. This construct was expressed in *E. coli* Rosetta 2 (DE3) cells. Cells were grown in 2xYT medium until OD600 reached 0.4 and induced with 0.4 mM IPTG for 4 hours at 37℃.

Cells were harvested by centrifugation, resuspended in lysis buffer (50 mM Tris pH 8.0, 300 mM NaCl, 0.5% Triton X-100 and 1 mM PMSF) and lysed by homogenisation. Lysates were centrifuged to remove cellular debris and protein purified by nickel-affinity purification using Ni-NTA resin (Qiagen) on a gravity flow column. Briefly, the supernatant containing the proteins was incubated with the resin for 30 minutes at 4℃ with nutation and applied to a column. The column was washed with 20 column volumes of wash buffer (50 mM Tris pH 8.0, 150 mM NaCl, 15 mM imidazole) and eluted with 5 column volumes of elution buffer (50 mM Tris pH 8.0, 150 mM NaCl, 250 mM imidazole). The eluted protein was concentrated and applied to Superdex S200 gel filtration column attached to AKTA Pure FPLC chromatography column (GE Healthcare) for observing co-elution.

##### Sequence Comparisons

The sequences for Vibrio parahaemolyticus VtrC (Q87GI3) and ToxS (Q05939), as well as the sequences for the known structures VpVtrA (5kewA) and VcToxR (7nn6A) and one PFAM(1) hit with borderline confidence (PsaF: WP_002216523) were used as queries for sensitive sequence search of structure (PDB_mmCIF70) and family (Pfam-A) databases provided by the HHPRED server(2, 3). Multiple sequence alignments (MSA) used for search were generated using PSI-BLAST against nr70, and all other parameters were set to default. Similar results were obtained using the default HHblits MSA generation on the HHPRED server.

Profiles corresponding to each VtrC-like family were built using the ProtBLAST/PSI-BLAST component of the HHpred server (5 iterations, 1e-3 E-value cutoff). The resulting multiple sequence alignments were converted to HMM profiles using HHmake from the HH-suite software package(4). The profiles were used to generate a database that was searched using each HMM as a query with HHsearch. HHsearch probabilities were reported for each query against all profiles in the database, with probabilities > 90 considered as confident.

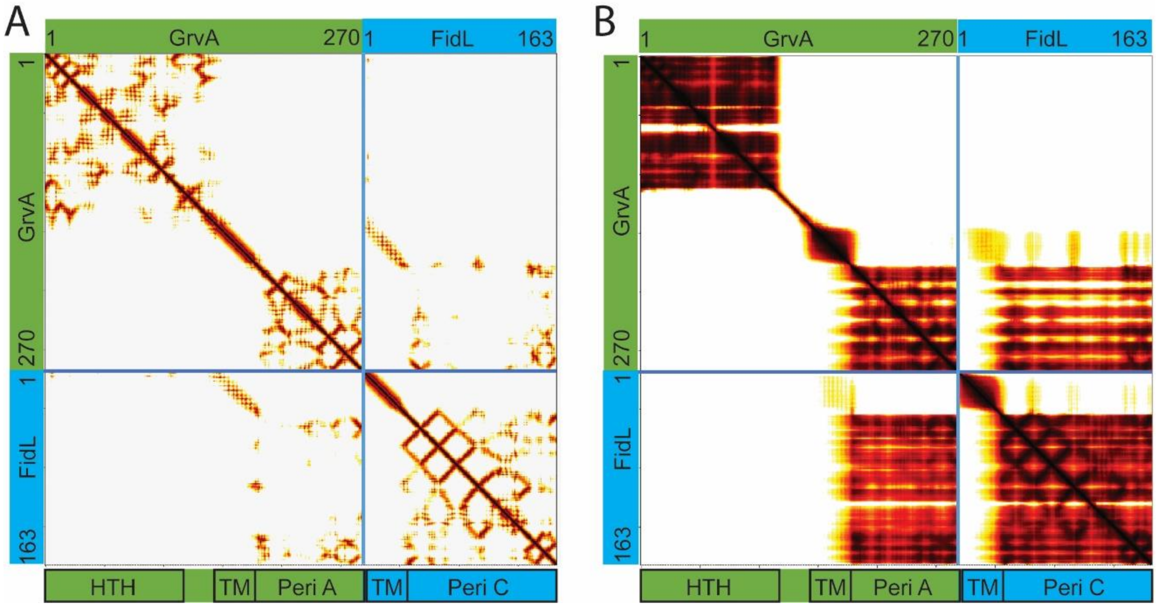


**Fig. S1. Residue distances and predicted aligned error for GrvA/FidL Structure Model**. Graphs represent residue numbers from the N-terminus to the C-terminus for the GrvA (green block) sequence, followed by the FidL (cyan block) sequence (labeled on both axes). The domain boundaries of GrvA (green) and FidL (cyan) are boxed and labeled below the graph. **A)** Observed distances between residue pairs, with darker colors indicating shorter distance. Pairs with distance > 12Å are not colored. **B)** Predicted aligned errors in distances between residue pairs, with darker colors indicating smaller predicted errors. Pairs with predicted error > 12Å are not colored.

```
VtrA           E  I  E  Y  *
             atgaaattgaatattaa
VtrC          M  K  L  N  I

TcpP          C  T  K  N  *
             atgcacaaaaaattaa
TcpH          M  H  K  K  L

YP3564        C  E  K  K  *
             atgcgaaaaaaaataa
YP3565        M  R  K  K  I

PsaE          C  K  Q  Q  *
             atgcaaacagcaatga
PsaF          M  Q  T  A  M

STM0341       C  I  T  V  K  E  R  Y  *
             atgtattacagtaaaagaacgttactga
STM0342       M  Y  Y  S  K  R  T  L  L

STM0344       C  I  S  I  K  E  R  Y  *
             atgcatttccataaaagaacgttattaa
STM0345       M  H  F  H  K  R  T  L  L

SF3508         D  E  T  Y  A  I  C  S  E  V  L  L  Y  D  *
             atgacgaaacttatgcaatttgttcagaggtgctattatatgactaa
SF3507        M  T  K  L  M  Q  F  V  Q  R  C  Y  Y  M  T

YP3283         E  M  V  N  C  *
             atgaaatggttaactgctag
YP3282        M  K  W  L  T  A

MarT          H  G  *
             catgggtaa
FidL          M  G
```

**Fig. S2. Co-component overlap of open reading frames**. Co-component receptor genes with overlapping open reading frames are labeled to the left of the corresponding encoded amino acid sequence. The upstream gene corresponding to the VtrA-like component is above the nucleotide sequence and the downstream gene corresponding to the VtrC-like component is below the nucleotide sequence. Overlapping Cys residues and their codons are colored red. Stop codons are indicated by *

**A**

| | **YqeI** | **GrvA** | **SF3508** | **STM0344** | **BprP** | ToxR | PsaE | YP3564 | YP3383 | VtrA | CadC | TcpP |
|---|---|---|---|---|---|---|---|---|---|---|---|---|
| **YqeI** | 1 | 0.666 | 0.499 | 0.533 | 0.458 | 0.361 | 0.362 | 0.296 | 0.242 | 0.232 | 0.181 | 0.136 |
| **GrvA** | 0.666 | 1 | 0.493 | 0.656 | 0.533 | 0.402 | 0.365 | 0.342 | 0.256 | 0.25 | 0.191 | 0.147 |
| **SF3508** | 0.499 | 0.493 | 1 | 0.468 | 0.443 | 0.284 | 0.359 | 0.274 | 0.24 | 0.217 | 0.192 | 0.191 |
| **STM0344** | 0.533 | 0.656 | 0.468 | 1 | 0.478 | 0.393 | 0.311 | 0.132 | 0.242 | 0.219 | 0.135 | 0.208 |
| **BprP** | 0.458 | 0.533 | 0.443 | 0.489 | 1 | 0.332 | 0.454 | 0.136 | 0.243 | 0.242 | 0.168 | 0.186 |
| ToxR | 0.361 | 0.402 | 0.284 | 0.393 | 0.332 | 1 | 0.319 | 0.294 | 0.397 | 0.325 | 0.25 | 0.181 |
| PsaE | 0.362 | 0.365 | 0.359 | 0.311 | 0.454 | 0.319 | 1 | 0.426 | 0.235 | 0.097 | 0.174 | 0.21 |
| YP3564 | 0.296 | 0.342 | 0.274 | 0.132 | 0.136 | 0.294 | 0.426 | 1 | 0.25 | 0.293 | 0.168 | 0.178 |
| YP3383 | 0.242 | 0.256 | 0.237 | 0.242 | 0.243 | 0.397 | 0.235 | 0.25 | 1 | 0.425 | 0.27 | 0.147 |
| VtrA | 0.232 | 0.25 | 0.217 | 0.219 | 0.246 | 0.325 | 0.097 | 0.293 | 0.425 | 1 | 0.283 | 0.174 |
| CadC | 0.181 | 0.191 | 0.192 | 0.126 | 0.168 | 0.25 | 0.174 | 0.168 | 0.27 | 0.283 | 1 | 0.152 |
| TcpP | 0.136 | 0.147 | 0.191 | 0.208 | 0.186 | 0.181 | 0.21 | 0.178 | 0.147 | 0.174 | 0.152 | 1 |

**B**

| | **YqeI** | **GrvA** | **SF3508** | **STM0344** | **BprP** | ToxR | PsaE | YP3564 | YP3383 | VtrA | CadC | TcpP |
|---|---|---|---|---|---|---|---|---|---|---|---|---|
| **YqeI** | 0 | 1.424 | 2.173 | 2.113 | 2.5 | 2.979 | 2.356 | 2.809 | 3.496 | 3.374 | 3.634 | 2.626 |
| **GrvA** | 1.424 | 0 | 1.929 | 2.053 | 1.806 | 2.474 | 2.631 | 2.198 | 3.241 | 3.41 | 3.361 | 3.153 |
| **SF3508** | 2.177 | 1.929 | 0 | 2.006 | 2.014 | 2.522 | 3.271 | 3.412 | 2.947 | 3.772 | 3.469 | 2.577 |
| **STM0344** | 2.113 | 2.053 | 2.006 | 0 | 2.188 | 3.416 | 2.101 | 3.008 | 2.977 | 3.242 | 3.057 | 3.163 |
| **BprP** | 2.5 | 1.904 | 2.148 | 2.188 | 0 | 2.571 | 1.642 | 3.437 | 3.027 | 3.249 | 3.098 | 3.924 |
| ToxR | 2.979 | 2.474 | 2.522 | 3.416 | 2.571 | 0 | 2.612 | 2.818 | 2.743 | 3.243 | 2.856 | 2.205 |
| PsaE | 2.359 | 2.631 | 3.271 | 2.101 | 1.642 | 2.612 | 0 | 1.958 | 2.256 | 2.607 | 2.411 | 1.77 |
| YP3564 | 2.809 | 2.198 | 3.412 | 3.008 | 3.437 | 2.818 | 1.958 | 0 | 2.734 | 3.07 | 3.132 | 2.343 |
| YP3383 | 3.496 | 3.241 | 2.947 | 3.112 | 3.027 | 2.743 | 2.256 | 2.734 | 0 | 2.432 | 3.273 | 3.736 |
| VtrA | 3.374 | 3.41 | 3.772 | 3.242 | 3.408 | 3.243 | 2.607 | 3.07 | 2.432 | 0 | 2.907 | 2.52 |
| CadC | 3.634 | 3.361 | 3.682 | 3.057 | 3.098 | 2.856 | 2.411 | 3.132 | 3.273 | 2.835 | 0 | 3 |
| TcpP | 2.626 | 3.153 | 2.577 | 3.163 | 3.924 | 2.205 | 1.77 | 2.343 | 3.736 | 2.52 | 3 | 0 |

**Figure S3. GESAMT pair-wise structure superposition heatmaps. A)** Pair-wise structure superpositions from GESAMT are colored in a Blue-white-red color scale from the highest (blue) to lowest (red) Q-score. **B)** Reported RMSDs are colored in the same scale, but from the lowest (blue) to highest (red). Structures are ordered the same as in Figure 5F from the main text.

**Table S1. HHpred Sequence Search**

| Query Gene | Sequence Range | Target ID | Target Database | Target Name | Target PDB | Prob-ability | Coverage |
|---|---|---|---|---|---|---|---|
| *VpVtrC* | 1-161 | 5KEW_B | PDB70 | VtrC | 5kewB | 100 | 0.83 |
| *VpVtrC* | 1-161 | PF17550.4 | PFAM | PsaF | na | 81.4* | 0.52 |
| *VpToxS* | 1-171 | PF17323.4 | PFAM | ToxS | na | 100 | 0.86 |
| *VpToxS* | 1-171 | PF18692.3 | PFAM | Duf6540 | 2m4l | 83.6* | 0.38 |
| *YpPsaF* | 1-162 | PF17550.4 | PFAM | PsaF | na | 100 | 1.00 |
| *YpPsaF* | 1-162 | PF13082.8 | PFAM | DUF3931 | na | 82.3* | 0.25 |
| *VcToxR* | 201-294 | 7nn6A | PDB70 | ToxR | 7nn6A | 100 | 0.89 |
| *VcToxR* | 201-294 | 6utcA | PDB70 | ToxR | 6utcA | 100 | 0.90 |
| *VcToxR* | 201-294 | 4osfA | PDB70 | Pan_kinase_C | 4osfA | 78.6* | 0.32 |
| *VpVtrA* | 165-263 | 5kewC | PDB70 | VtrA | 5kewC | 100 | 1.00 |
| *VpVtrA* | 165-263 | PF08350.12 | PFAM | Duf1724 | na | 55.6* | 0.41 |

*Unconfident HHpred Probability < 90

**Table S2.** Reference Genomes.

| Organism | Strain | Organism Groups | Assembly |
|---|---|---|---|
| *Mycobacterium tuberculosis* | H37Rv | Actinobacteria | GCF_000195955.2 |
| *Chlamydia trachomatis* | D/UW-3/CX | Chlamydiae | GCF_000008725.1 |
| *Staphylococcus aureus* subsp. aureus | NCTC 8325 | Firmicutes | GCF_000013425.1 |
| *Listeria monocytogenes* | EGD-e | Firmicutes | GCF_000196035.1 |
| *Bacillus subtilis* subsp. Subtilis | 168 | Firmicutes | GCF_000009045.1 |
| *Caulobacter vibrioides* | NA1000 | Alphaproteobacteria | GCF_000022005.1 |
| *Campylobacter jejuni* subsp. jejuni | NCTC 11168 | delta/epsilon | GCF_000009085.1 |
| *Acinetobacter pittii* | PHEA-2 | Gammaproteobacteria | GCF_000191145.1 |
| *Coxiella burnetii* | RSA 493 | Gammaproteobacteria | GCF_000007765.2 |
| *Escherichia coli* | K-12 substr. MG1655 | Gammaproteobacteria | GCF_000005845.2 |
| *Escherichia coli* O157:H7 | Sakai substr. RIMD 0509952 | Gammaproteobacteria | GCF_000008865.2 |
| *Klebsiella pneumoniae* subsp. Pneumoniae | HS11286 | Gammaproteobacteria | GCF_000240185.2 |
| *Pseudomonas aeruginosa* | PAO1 | Gammaproteobacteria | GCF_000006765.1 |
| *Salmonella enterica* subsp. enterica serovar Typhimurium | LT2 | Gammaproteobacteria | GCF_000006945.2 |
| *Shigella flexneri* | 301 | Gammaproteobacteria | GCF_000006925.2 |
| *Vibrio cholerae* | MS6 | Gammaproteobacteria | GCF_000829215.1 |
| *Vibrio parahaemolyticus* | O3:K6 substr. RIMD 2210633 | Gammaproteobacteria | GCF_000196095.1 |
| *Yersinia pseudotuberculosis* | IP32953 | Gammaproteobacteria | GCF_000834295.1 |

**Table S3** VtrC-like Component HHsearch Comparisons

| | FidL | YqeJ | BprQ | Sf3507 | STM 0345 | STM 0432 | Yp3565 | TcpH | ToxS | Yp3282 | PsaF | VtrC |
|---|---|---|---|---|---|---|---|---|---|---|---|---|
| FidL | 100 | 100 | 99.9 | 97 | 97 | 56.4 | 62 | 1.4 | 51.8 | 0.6 | 43.2 | 5.3 |
| YqeJ | 100 | 100 | 99.6 | 96.3 | 83.1 | 33.8 | 78.7 | 0.2 | 10 | 0.6 | 17.4 | 2.6 |
| BprQ | 99.9 | 99.6 | 100 | 97.7 | 97.4 | 4.2 | 85.2 | 0.1 | 1.4 | 3.6 | 10.9 | 1.2 |
| Sf3507 | 97.2 | 96.6 | 98 | 100 | 1.3 | 6.4 | 0.2 | 0.3 | 0.5 | 0.3 | 0.5 | 7.7 |
| STM0345 | 97.3 | 86 | 97.7 | 1.4 | 100 | 100 | 0.1 | 0.1 | 0.2 | 0.2 | 0.1 | 0.9 |
| STM0432 | 63.6 | 38.1 | 6.7 | 6.7 | 100 | 100 | 0.1 | 0.1 | 1.9 | 0.2 | 0.1 | 0.1 |
| Yp3565 | 67.3 | 81.2 | 88.7 | 0.3 | 0.1 | 0.2 | 100 | 0.1 | 0.6 | 0.1 | 0.1 | 0.1 |
| TcpH | 1.4 | 0.2 | 0.2 | 0.3 | 0.1 | 0.1 | 0.1 | 100 | 0.1 | 0.2 | 0.1 | 0 |
| ToxS | 56 | 8.6 | 1.6 | 0.4 | 0.1 | 1.4 | 0.3 | 0 | 100 | 0.2 | 0.7 | 1.5 |
| Yp3282 | 0.6 | 0.5 | 5.6 | 0.3 | 0.2 | 0.2 | 0.1 | 0.2 | 0.3 | 100 | 0.1 | 0.2 |
| PsaF | 43.1 | 16 | 15.1 | 0.5 | 0.1 | 0.1 | 0.1 | 0.1 | 1 | 0.1 | 100 | 30.1 |
| VtrC | 5.3 | 2.4 | 1.9 | 6.9 | 0.9 | 0.2 | 0.1 | 0 | 2 | 0.3 | 28.5 | 100 |

Confident HHsearch Probability > 90 is highlighted gray

**SI References**


1. J. Mistry *et al.*, Pfam: The protein families database in 2021. *Nucleic Acids Res* **49**, D412-D419 (2021).
2. J. Soding, A. Biegert, A. N. Lupas, The HHpred interactive server for protein homology detection and structure prediction. *Nucleic Acids Res* **33**, W244-248 (2005).
3. L. Zimmermann *et al.*, A Completely Reimplemented MPI Bioinformatics Toolkit with a New HHpred Server at its Core. *J Mol Biol* **430**, 2237-2243 (2018).
4. M. Steinegger *et al.*, HH-suite3 for fast remote homology detection and deep protein annotation. *BMC Bioinformatics* **20**, 473 (2019).
